# The brain architecture of punishment learning

**DOI:** 10.1101/2025.11.24.690287

**Authors:** Alexandra V. Gregory, Philip Jean-Richard-dit-Bressel, Nikki Huang, James Diefenbach, EunA Choi, Jason Soo, Jessica Chen, A. Simon Killcross, Gavan P. McNally

## Abstract

Learning which actions cause harm is essential for survival, yet how the brain transforms this knowledge into adaptive control of behavior remains unclear. We combined whole-brain analysis, multisite chemogenetic silencing, spatial transcriptomics, and longitudinal calcium imaging to map how punishment learning reorganizes brain networks across scales. Learning reshaped mesoscale community organization into a network anchored by the amygdala, subthalamic–hypothalamic zone, and ventral midbrain tegmentum. These regions contributed distinct components of adaptive avoidance and engaged diverse transcriptional programs across multiple neuronal subclasses. Within this network, the amygdala acted as a hub, partitioning action and outcome information and remapping neural geometry to segregate punished from safe actions. Together, these results identify a multiscale neural architecture through which aversive experience can be transformed into flexible behavioral control.

---

Learning to modify behavior in response to aversive consequences is essential for survival. Despite this importance, the neural mechanisms by which punishment shapes our behaviors and choices remain poorly understood. Much of what is known derives from Pavlovian fear paradigms, where environmental cues acquire threat value and elicit involuntary defensive responses (*1*, *2*). These accounts provide limited insight into how organisms voluntarily regulate their own actions when the actions themselves cause harm, a core feature of adaptive behavior.

Punishment learning poses a distinct challenge. Rather than linking external cues to threat, it requires the brain to learn relationships between self-generated actions and aversive outcomes, and to adjust behavior by selectively suppressing the dangerous action while continuing to pursue reward through alternative means (*3*, *4*). Influential reinforcement learning models treat this process as the negative of reward: a scalar inversion of value that reduces response strength (*5–8*). Yet punishment learning depends on more complex cognitive process than just valuation (*9–11*). The brain mechanisms that subserve this learning to enable precise, response-selective suppression are largely unknown (*12*). How does the brain transform the representation of an action once it becomes associated with harm?

Learning and action control arise from distributed brain networks and activity across neural scales (*12–16*). To capture this multiscale organization, we used unbiased whole-brain analysis, multisite chemogenetics, spatial transcriptomics, and longitudinal calcium imaging to map the brain architecture of punishment learning. This analysis showed that punishment learning preserves the brain’s small world topology while reconfiguring its large-scale functional organization. Within this reconfigured network, several hub regions emerged whose activity was associated with complementary facets of adaptive avoidance and whose transcriptional programs revealed a diverse set of neuronal subclasses engaged by punishment learning. Among these, the amygdala served as an integrative hub whose ensembles dynamically transformed action representations, recoding punished actions into an aversive neural space. Together, these findings identify a multiscale architecture supporting the transformation of aversive experience into adaptive behavioral avoidance.

## Re-organization of brain networks during punishment learning

We first established the behavioral effects of punishment using a within-subjects, two-response task (**Fig 1A**). In the punishment condition, one response (R1) delivered food reward on a variable interval (VI) schedule but also delivered a footshock punisher under a fixed ratio (FR8) schedule, whereas the alternative response (R2) delivered food without shock. Control animals experienced the same food contingencies in the absence of punishment. Adaptive avoidance was evident as selective suppression of the punished response (R1) relative to the unpunished response (R2), which increased steadily over training blocks compared to controls (**Table S7** reports all statistical analyses). This response-selective suppression is the behavioral signature of punishment learning (*3*, *10*). To confirm that suppression was due to the response-punisher (i.e. instrumental) contingency and not a stimulus - punisher (i.e. Pavlovian) contingency we tested a separate group of mice using a within-subjects yoking task (**Fig S1**). Here, R1 again delivered food reward on a VI schedule and a footshock punisher under an FR8 schedule. We recorded the timing of punisher delivery and then played these back in a response non-contingent manner to the same animal during access to R2. This within-subjects yoking matches R1 and R2 on Pavlovian contingencies but separates them on instrumental contingency. Only the instrumental contingency suppressed lever pressing.

**Fig. 1.**
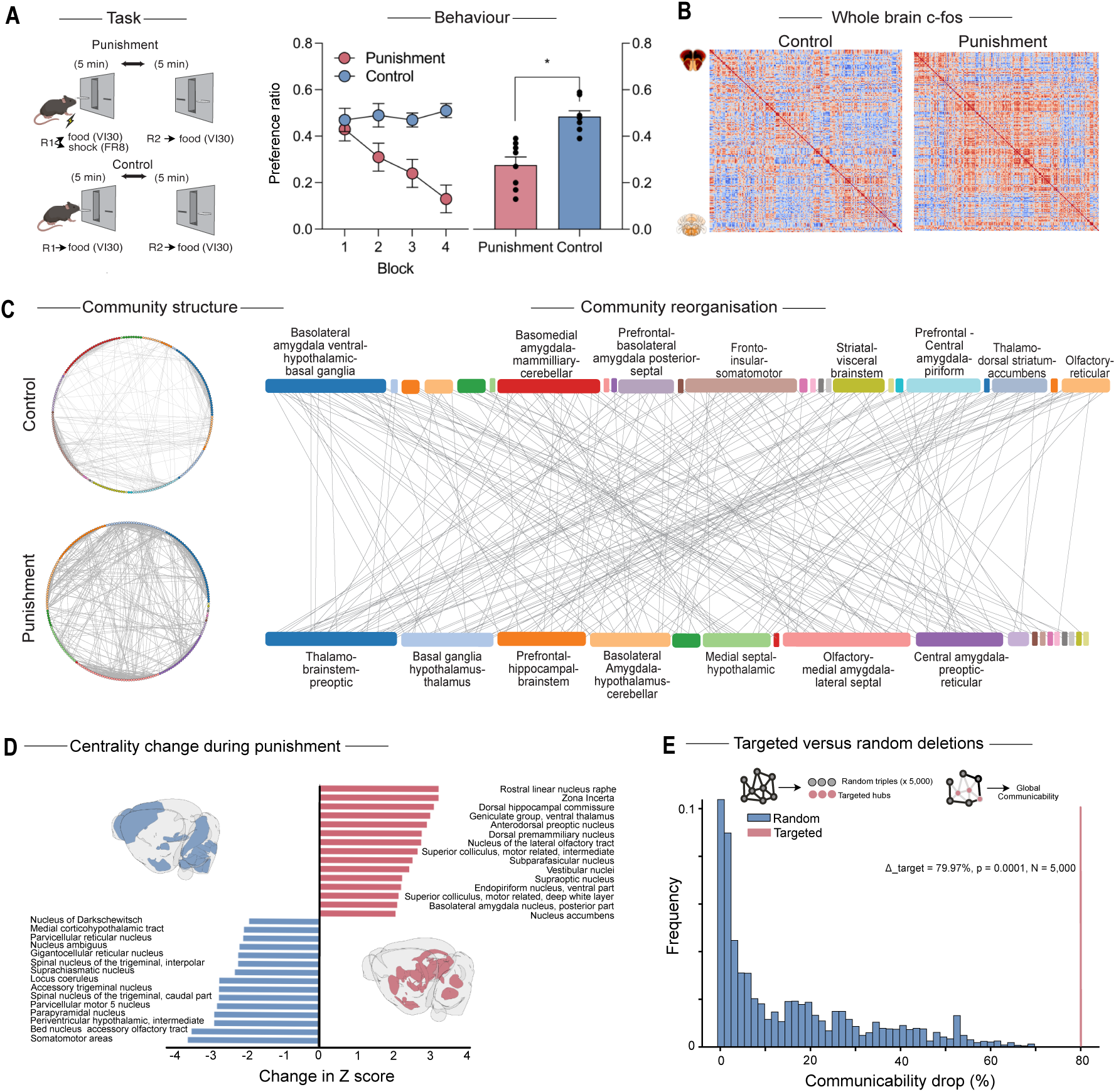
Network analysis of whole-brain c-Fos identifies a distinct punishment network. (A) Experimental design. Punished mice (n = 8) were trained with R1 → punishment (0.3 mA, 0.5 s) + reward and R2 → reward in alternating five-minute blocks. Control mice (n = 8) received R1 → reward and R2 → reward. Mean ± SEM preference ratios are shown across five-minute training blocks and averaged across the session. (B) Functional connectivity. Whole-brain c-Fos Pearson correlation matrices for Punished and Control groups. (C) Community organization. Circle plots illustrating the community structure of brain networks in Punished and Control groups. Each color represents a distinct community of regions that co-activate. The tanglegram links the same brain regions across groups, highlighting how community memberships reorganize following punishment learning. (D) Network centrality. Change in normalized degree centrality for the top and bottom 15 brain regions. (E) Network robustness. In silico targeted versus random node deletion on network communicability. Statistical analysis: p < 0.05 independent-means t-test (A) and permutation test (E).

Next, we asked how punishment learning is organized in the brain. To map brain-wide reorganization accompanying punishment learning (**Fig 1A**), we combined tissue clearing, immunolabelling, and light-sheet imaging to quantify expression of the activity marker Fos across 206 regions (**Fig 1, Fig S2**). Treating brain regions as nodes and inter-regional correlations as edges (*16*, *17*), we tested whether punishment alters large-scale organization (*18–20*). Correlation matrices of Fos counts (**Fig 1B**) were used to generate co-activation networks (**Fig S3**). Both Control and Punishment networks exhibited modular, small-world topology (**Fig S3**), indicating preservation of global principles of brain networks (*18*, *19*, *21*). However, community structure differed markedly between groups (**Fig 1C, Tables S1-S2**) and similarity (Jaccard, adjusted rand index, and normalized mutual information) was low (**Table S3**), showing that punishment learning operates through large-scale reorganization, reallocating regional participation across modules rather than merely changing connection strength within fixed communities. Notably, under punishment, a community emerged linking prefrontal regions (anterior cingulate, prelimbic, infralimbic, orbital cortices), limbic (hippocampal), and brainstem neuromodulatory structures (superior colliculus, locus coeruleus, laterodorsal tegmental nucleus), consistent with roles in decision-making (*22*, *23*), action control (*24–26*), and emotional state regulation (*27*, *28*).

We compared Punishment and Control networks to identify regions whose relative influence within the functional architecture changed with learning (**Fig 1D**). Increases in centrality indicate enhanced integration or connectivity, whereas decreases reflect reduced participation in network communication. Regions involved in behavioral automaticity (e.g., somatomotor cortex) (*24*, *25*) diminished in importance during punishment, whereas others across the amygdala, subthalamic–hypothalamic zone, and midbrain increased, including the basolateral amygdala (BLA), zona incerta (ZI), and rostral linear nucleus of the raphe (RLi) (Figure 1D). These regions emerged as hub nodes during punishment, consistent with their roles in learning (*29–34*), valence (*33*, *35–39*), and sensory–motor integration (*40–42*). *In silico* deletion of these three hubs from the punishment network, individually or together, had minimal effects on basic network metrics (number of nodes/edges, direct connections, shortest paths) whereas triple but not individual deletion disrupted global communicability, a measure of how efficiently signals can travel across the entire network through both direct and indirect pathways (**Fig S3**). To test specificity, we generated a null distribution by deleting 5,000 random triples of nodes from the Punishment network. Most random deletions produced only modest reductions in communicability, reflecting the redundancy of most regions. In contrast, simultaneous removal of the BLA, ZI, and RLi caused a marked reduction in global communicability, far outside the null distribution **(Fig 1E**). Thus, these hub nodes are critical for maintaining network integration under punishment conditions. These *in silico* analyses generate a causal prediction: if punishment learning depends on reallocating these brain regions network participation, then silencing these hubs should impair adaptive, response-selective suppression.

## Parallel processing during punishment learning

To test this network-level prediction and bridge computational inference and causal validation, we used multisite chemogenetic inhibition during punishment learning. Mice received infusions of the inhibitory DREADD AAV5-hSyn-hM4Di into the amygdala (targeting BLA), subthalamic–hypothalamic zone (targeting ZI), and ventral midbrain tegmentum (targeting RLi) concurrently, with controls receiving AAV5-hSyn-eGFP (**Fig 2A, Fig S4**). Mice were trained in the same within-subjects punishment task as before. Prior to punishment training, they received injections of the DREADD ligand CNO (3 mg/kg) (**Fig 2B**) or vehicle (**Fig S5**) and were tested 48 hr later in the absence of punishment.

**Fig. 2.**
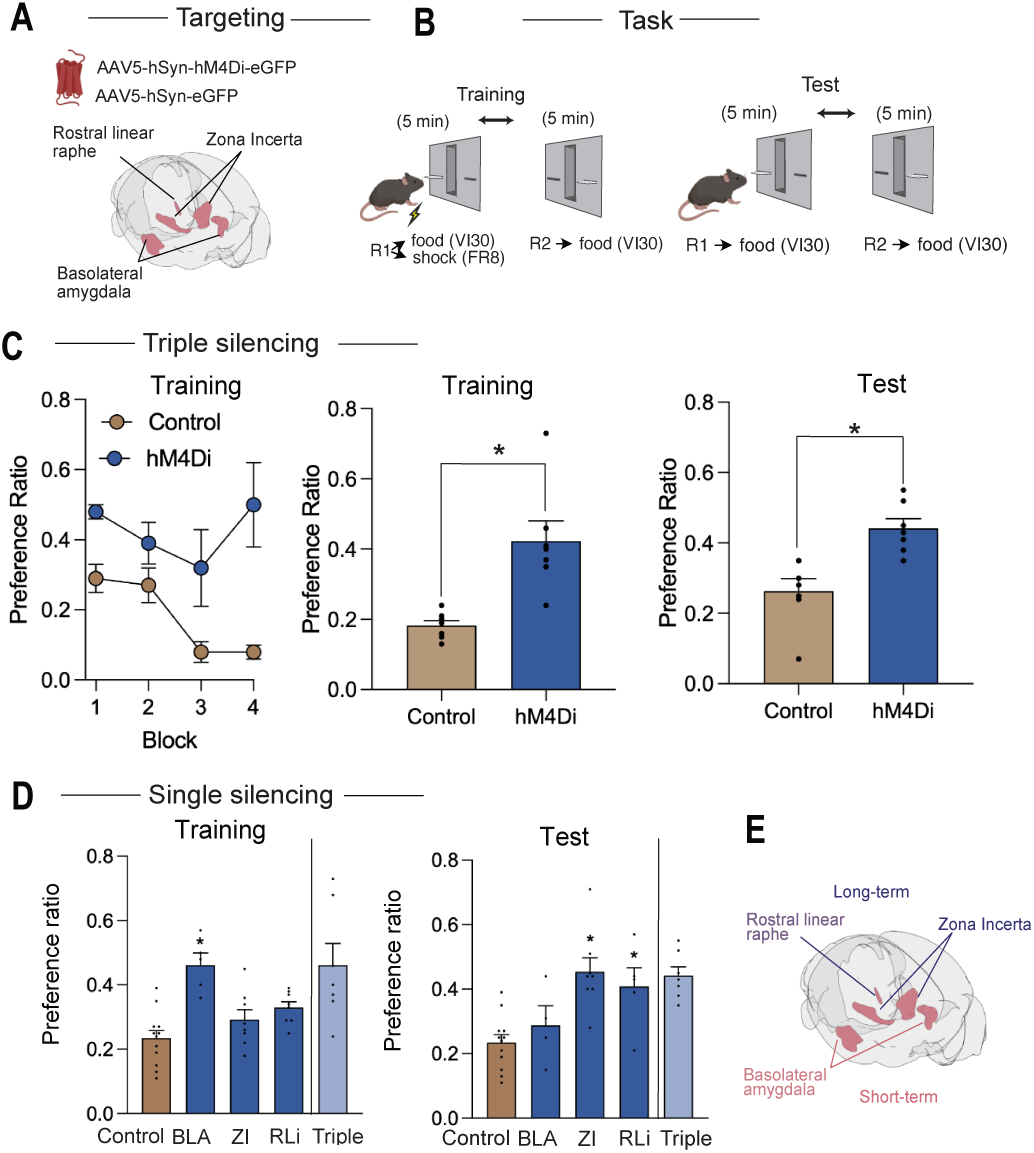
Chemogenetic silencing identifies distinct hub-node roles in adaptation to punishment. (A) Viral targeting. Mice received AAVs encoding the inhibitory DREADD hM4Di or control eGFP in the basolateral amygdala (BLA), zona incerta (ZI), and rostral linear nucleus (RLi). (B) Experimental design. Mice were trained with R1 → punishment (0.3 mA, 0.5 s) + reward and R2 → reward in alternating five-minute blocks. Intraperitoneal injections of CNO (3 mg/kg) or vehicle (see Fig S5) were administered before training, and testing occurred 48 h later. (C) Triple-region silencing. Preference ratios across five-minute training blocks, overall training, and test sessions (hM4Di n = 7; Control n = 7). (D) Single-region silencing. Training and test preference ratios for each region (triple-silencing data shown for comparison) (Control n = 12; BLA n = 5; ZI n = 8; RLi n = 8). (E) Summary. Single-region silencing effects. Statistical analysis: p < 0.05, independent-means t-test (C); two-way followed by one-way ANOVA and pairwise comparisons against Control (D). Data are mean ± SEM.

In controls expressing eGFP, punishment produced selective suppression of R1 relative to R2 across training blocks, diagnostic of punishment learning (**Fig 2C**). As anticipated from the network analysis, concurrent hM4Di inhibition of amygdala, subthalamic–hypothalamic zone, and ventral midbrain tegmentum profoundly impaired this learning. Mice failed to show normal selective suppression of the punished response during training and test (**Fig 2C**). This behavioral result mirrors the *in silico* deletion results, where simultaneous removal of these hubs produced a collapse in network communicability, and demonstrates that these brain regions are required for punishment learning *in vivo*. hM4Di inhibition did not affect latency to lever press, reward seeking, or the number of punishers received (**Fig S5**), confirming that effects were selective to learning rather than to general performance. Inspection of lever-pressing patterns further showed that hM4Di inhibition prevented selective association of the punished response with punisher delivery, yielding generalized suppression of lever pressing in the hM4Di but not the eGFP group (**Fig S5**). So, punisher efficacy was intact; the chemogenetic deficit reflects impaired response-selective association rather than sensory or motivational loss. This confirms that these brain regions are necessary for adaptive punishment avoidance rather than nonspecific behavioral inhibition.

We next asked whether any single hub could account for this effect. Using the same hM4Di inhibition and behavioral training approach, we targeted the amygdala, subthalamic–hypothalamic zone, and ventral midbrain tegmentum individually. Chemogenetic silencing centered on BLA impaired within-session (*34*, *43*) but not between-session adaptation. Silencing centered on ZI spared within-session but impaired between-session adaptation **(Fig 2D, Fig S5**). Silencing centered on RLi largely spared within-session (there was a small effect under either CNO or vehicle, indicating non-specific effects of hM4Di transduction) and impaired between-session adaptation.

These behavioral findings refine the network prediction, showing that amygdala, subthalamic–hypothalamic zone, and ventral midbrain tegmentum make complementary, not redundant, contributions to punishment learning. Amygdala activity underpinned within session adaptation to punishment contingencies whereas subthalamic–hypothalamic and midbrain activity bridged this to longer-term, between-session adaptation. Thus, parallel processes across the distributed punishment network support adaptive behavioral avoidance (**Fig 2E**).

## Transcriptional architecture of punishment learning

Having identified amygdala, subthalamic–hypothalamic zone, and ventral midbrain tegmentum as key hubs in the punishment network, we next investigated the cell types associated with these roles. We profiled spatial transcriptomes across these regions to identify how transcriptional activity is organized during punishment learning. To do this, mice (n = 4, Punished, n = 4 Control) were trained in the within-subjects punishment task (**Fig 3A,B**) and the Vizgen MERSCOPE platform was used to profile spatial gene expression across coronal sections containing the amygdala, subthalamic–hypothalamic zone, and ventral midbrain tegmentum (**Fig 3C**). We applied consensus non-negative matrix factorization to identify latent gene expression programs recurring across cells from all animals in these regions (*44*). Similarity analyses and leave-one-out resampling validated punishment-specific programs in each region (**Table S4**) and MapMyCells was used to catalogue cell sub-classes expressing these gene programs as their top program (**Fig 3D, Table S5, S6)**.

**Fig. 3.**
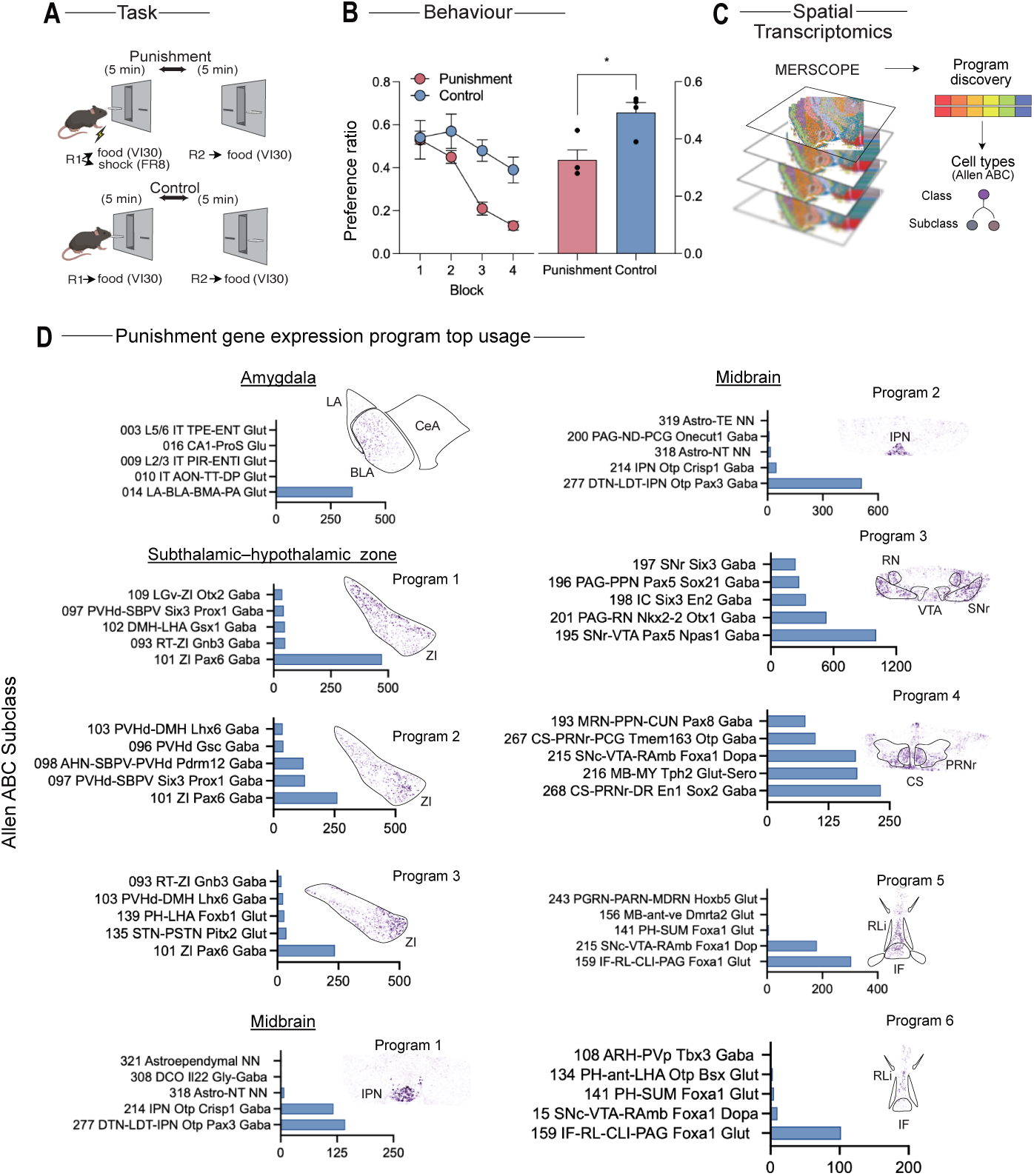
Spatial transcriptomics identifies region- and cell-type-specific gene-expression programs engaged by punishment learning. (A) Task. Mice (n = 4) were trained with R1 → punishment (0.3 mA, 0.5 s) + reward and R2 → reward in alternating five-minute blocks. Control mice (n = 4) received R1 →reward and R2 → reward. (B) Behavior. Mean ± SEM preference ratios across five-minute training blocks and averaged across the session (note: 2 animals in punished group had preference ratios = 0.3). (C) Spatial transcriptomics workflow. Tissue sections from the amygdala, subthalamic–hypothalamic zone, and midbrain were imaged using the MERSCOPE platform. Gene-expression programs were derived by consensus non-negative matrix factorization (cNMF) and mapped to cell types defined by the Allen Brain Cell (ABC) Atlas. (D) Punishment-specific gene-expression programs. Top program usage by brain region and Allen ABC subclass. Coronal sections show heatmaps of top program usage. Statistical analysis: p < 0.05, repeated-measures ANOVA (B). Data are mean ± SEM (B) and cell counts (D).

This analysis revealed a heterogeneous molecular architecture across the punishment network. In the amygdala, a program was detected in BLA glutamate neurons whose top genes related to plasticity (e.g., *Bdnf*), neuropeptide binding (e.g, *Cckbr, Sstr2, Oprd1*) and synaptic organization (e.g., *Ntng2*). Three programs, with subtly different spatial locations, were detected in the ZI, predominantly in Pax6-GABA neurons implicated in multisensory integration, anxiety and motivational control (*31*, *40*, *45–47*). Multiple distinct punishment programs were detected in the ventral midbrain tegmentum. Midbrain programs 1 and 2, centered in the interpeduncular nucleus (IPN), comprised Otp-Pax3 and Otp-Crisp1 GABAergic populations implicated in amplifying responses to aversive events (*48*). Programs 3 and 4 involved Pax5, Otx1, and En1 GABA as well as Foxa1 dopamine neurons spanning the ventral tegmental area, substantia nigra reticulata (SNr), red nucleus (RN), pontine reticular nucleus (PRN), regions associated with action selection and motor control (*15*, *49*, *50*). Programs 5 and 6 centered on RLi and encompassed Foxa1 glutamatergic and dopaminergic neurons (*49*, *50*) linked to mesocorticolimbic and mesothalamic pathways (*51*) as well as aversion signaling (*37*, *52*)

These findings show that distinct cell type–specific transcriptional adaptations occur within the amygdala, subthalamic–hypothalamic zone, and ventral midbrain tegmentum in punishment. The general neuronal populations identified align well with known roles in aversion, motivational control, and behavioral flexibility (*30*, *37*, *50*): but these results considerably refine that understanding by identifying specific cell sub-classes engaged and expanding the catalogue of cells associated with punishment learning.

## Stable and flexible ensemble coding during punishment learning

Our whole brain network and spatial transcriptomic analyses present static, end-point measures of neural activity. They do not show which events are critical drivers of this activity or how activity dynamically unfolds across longer timescales. To address this, we turned to longitudinal *in vivo* calcium imaging to track neuronal ensemble dynamics. We expressed GCaMP7f in BLA neurons and implanted a GRIN lens to record these neurons using miniature head-mounted fluorescent microscope (*53*) (**Fig 4A, Fig S7**). We focused on BLA because of its status as a hub node in the brain network for punishment (Figure 1, 2) (*33*, *34*, *43*), its broad reciprocal connectivity with the cortex (*54–56*), and the recruitment of plasticity-related gene programs by punishment learning (**Fig 3**). Mice were trained in a multi-session variant of the within-subjects punishment task. Across six days of punishment training, they learned to selectively suppress the punished response (R1). During a choice test, when R1 and R2 were available concurrently in the absence of punishment, mice preferred the safe lever (R2) over the previously punished lever (R1). Subsequently, across three days of extinction training in which the punisher was omitted, mice flexibly updated their learning and selectively increased responding on R1 (**Fig 4B**).

**Fig. 4.**
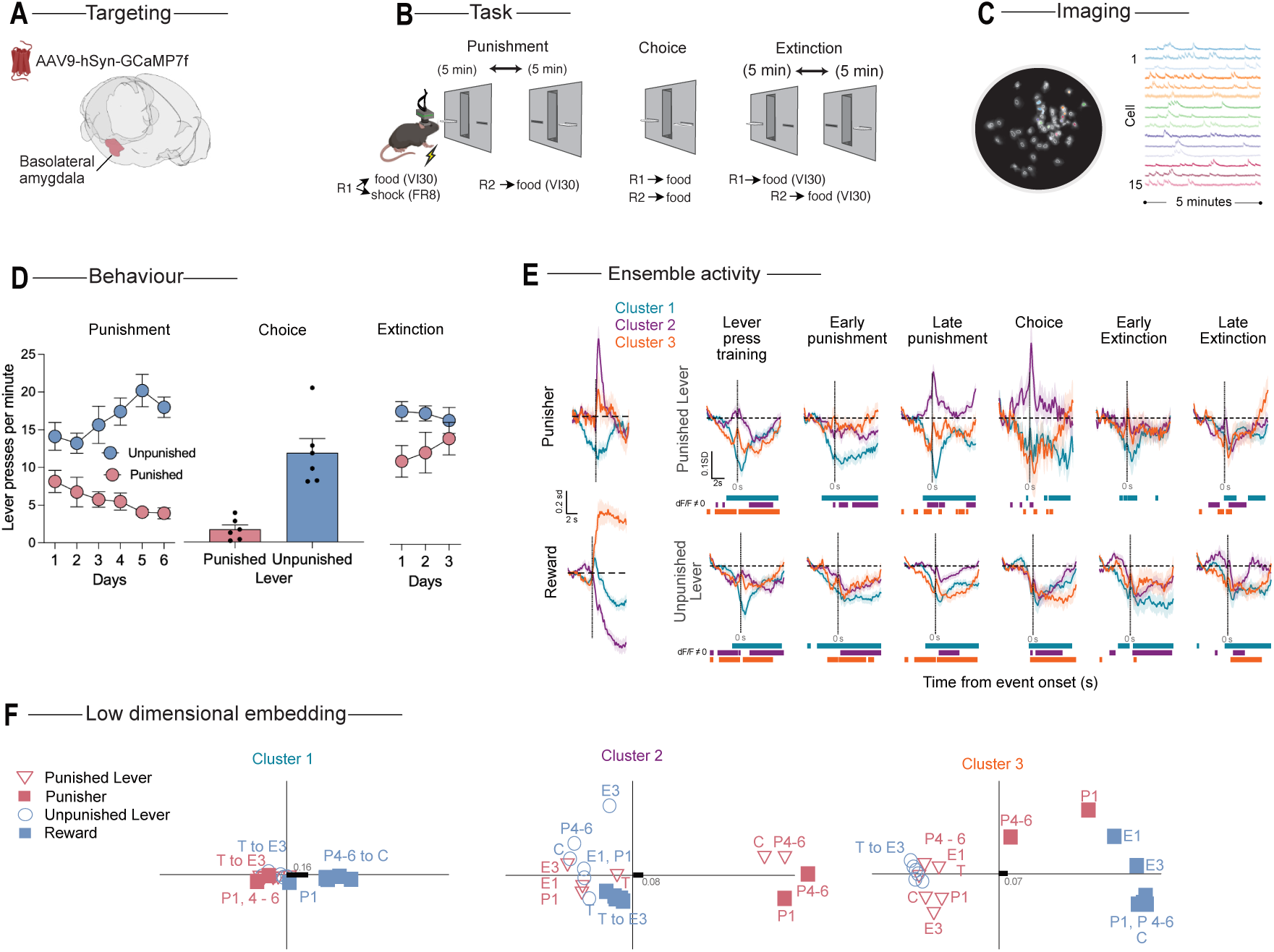
Longitudinal in vivo calcium imaging identifies neuronal-ensemble dynamics during punishment learning. (A) Targeting. AAV9-hSyn-GCaMP7f was injected into the basolateral amygdala for calcium imaging of neuronal activity, and mice received GRIN-lens implantation. (B) Task. Mice (n = 6) were trained with R1 → punishment + reward and R2 → reward in alternating five-minute blocks, followed by choice and extinction tests. (C) Imaging. Representative miniscope field of view showing individual identified cells (left) and raw calcium traces (right). (D) Behavior. Lever presses per minute (mean ± SEM) across punishment, choice, and extinction phases. (E) Longitudinal ensemble activity. Cluster-based kernels for each event category of interest at each experimental phase. Significant cluster-based kernel transients within the −5 to 5 s peri-event period were identified via 95% bootstrapped confidence intervals and are shown by colored lines below each plot. Cluster 1: 165 cells; Cluster 2: 270 cells; Cluster 3: 176 cells. (F) Low-dimensional embedding. Multidimensional scaling of normalized fit scores for peri-event kernels showing the evolution of ensemble geometry; black bars show metric stress. Statistical analysis: p < 0.05, repeated-measures ANOVA (D).

We recorded Ca²⁺ activity from 1,147 BLA neurons in 6 freely moving mice (**Fig 4C**) allowing us to capture how BLA neuron activity evolved as animals learned punishment, used that learning to guide choice, and flexibly updated behavior when contingencies changed (**Fig 4D**). As expected (*57*), not all BLA neurons were outcome-responsive and they responded via excitatory and inhibitory transients. The distribution of these responses changed across punishment training. More BLA neurons were responsive to punishment and reward across punishment training (**Fig S7**). We assessed ensemble-level structure in BLA activity (**Fig 4E**). To do this, we longitudinally tracked neurons across nine sessions (lever-press training [Days 9 –10]; early punishment [Day 1], Late punishment [Days 4–6]; choice; and extinction [Days E1–3)] (**Fig 4B,D**). Unbiased, data-driven clustering of normalized peri-event transients identified three distinct clusters (**Fig 4E**). Cluster 1 (n = 165) neurons were inhibited by the punisher, showed a biphasic excitatory–inhibitory response to reward, and were inhibited during lever pressing. Cluster 2 (n = 270) neurons were excited by the punisher but inhibited by reward. Cluster 2 neurons were always inhibited during unpunished lever pressing but developed excitatory responses to the punished lever across punishment sessions that reverted back to inhibitory responses during extinction, reflecting changes in action value and avoidance across these phases. Cluster 3 (n = 176) neurons exhibited an immediate excitatory response to punishment, a delayed excitatory response to reward, and inhibition during lever pressing. These findings show a dynamic reorganization of BLA ensemble coding during punishment learning, with specific neuronal populations differentially tracking punishment and reward as well as the actions that produce them.

To determine how these clusters relate to behavior across sessions and whether their activity reorganizes across learning, we quantified similarity among cluster-based peri-event kernels and visualized their geometry in a low dimensional space using multidimensional scaling (*58*, *59*). Similarity between kernel waveforms was quantified with 95% confidence intervals for fits derived from the 2.5th and 97.5th percentiles of bootstrapped distributions (1,000 random resamples of mean transients) (*60*). Waveforms were deemed significantly similar if their fit confidence interval lay entirely above zero, or significantly inverse if it lay entirely below zero (*60*).

This analysis showed a tripartite organization (**Fig 4F, Fig S7**). Cluster 1 neurons stably discriminated reward from all other events across training, punishment, and extinction. Cluster 2 neurons exhibited response-selective and contingency-sensitive remapping. Prior to and early in punishment training, Cluster 2 neuron punished lever activity was similar to reward; as punishment learning progressed, this activity became dissimilar to reward and aligned with punishers. When the punisher was omitted during extinction, activity reverted to the reward pattern. Cluster 3 neurons partitioned actions from outcomes, stably discriminating lever presses from punishment and reward, but not differentiating within either of these categories. Critically, cluster activity selectively predicted emergence of adaptive punishment avoidance (**Fig S7**), linking ensemble dynamics to response-selective suppression.

Together, these analyses show that longitudinal BLA ensemble activity mirrors the associative and evolving behavioral structure of punishment learning, encoding stable representations of reward (Cluster 1), maintaining a consistent partition between actions (antecedents) and outcomes (consequences) (Cluster 3), and flexibly adapting response-specific activity patterns to represent the contingencies of punished actions (Cluster 2). Thus, BLA ensembles exhibit both stability and flexibility, providing a substrate for response-selective punishment learning.

## Discussion

Adaptive behavior requires learning which actions yield reward and which lead to harm. Actions that cause harm must be selectively suppressed without abolishing the capacity to act and pursue reward. Our findings outline a multiscale neural architecture through which aversive experience can be transformed into this flexible behavioral control

At the macroscale, punishment learning reconfigures the brain’s large-scale community structure. The global small-world and modular organization of brain networks (*18*) was preserved, but the composition of modules and the regional affiliations within them, shifted markedly during punishment learning. This dynamic reallocation of regional participation shows that punishment learning follows the same principle of flexible network topology described for task switching and cognitive control (*19*, *61–63*). At the mesoscale, distributed regions contributed complementary functions for transforming immediate aversive experience into lasting, adaptive control. BLA enabled rapid, within-session adjustment; ZI and RLi supported longer-term, between-session adjustment revealing distinct substrates for working memory and long-term consolidation of punishment learning. At the microscale, punishment learning recruited a broad, diverse cellular repertoire and region-specific transcriptional programs. More than 17 subclasses were identified across key nodes of the network, including BLA glutamate neurons involved in valence coding (*30*, *36*, *64*) and region-specific GABAergic subclasses in ZI and RLi/midbrain implicated in aversion and behavioral flexibility (*31*, *37*, *40*, *45–47*, *50*, *52*).

Within this multiscale architecture, BLA emerged as a hub for adaptive control. Its ensembles completed a minimal representational basis for instrumental learning: one ensemble stably encoded reward; a second maintained a consistent partition between actions (antecedents) and outcomes (consequences); a third flexibly remapped response-specific activity to represent punished actions. This organization offers a plausible mechanistic basis for BLA’s contribution to flexible behavior. During learning, representations of punished actions shifted toward an aversive coding space, consistent with observed behavioral inhibition of the action, whereas unpunished actions remained aligned with reward. Similar population realignments occur in instrumental reward learning (contingency/value shifts (*65*, *66*)) and Pavlovian fear (CS convergence on US; (*57*)), suggesting a shared principle whereby learning transforms predictors, cues or actions, into the outcome’s representational space within BLA topology (*11*). In punishment, this transformation is instantiated within an action–outcome space, linking internal decisions to aversive outcomes.

These findings situate punishment learning within a broader understanding of adaptive control, in which brain regions flexibly assemble into a distributed network, characterized by diverse cell subclasses and ensemble remapping, to address fundamental adaptive challenges. Impairments in punishment learning are recognized as transdiagnostic features of psychopathology, underlying failures to suppress harmful or self-destructive behaviors in conditions such as addiction, obsessive–compulsive disorder, and psychopathy (*67*, *68*). The mechanisms identified here suggest a multiscale substrate for adaptive control. Disruption of these processes at any level could contribute to the impaired decision-making observed across disorders involving punishment insensitivity.

## Acknowledgments

We thank Iveta Slaptova and the Katarina Gaus Light Microscopy Facility (UNSW) for assistance with MERSCOPE. The following AI-assisted technologies were used in preparation of the manuscript: OpenAI ChatGPT4 and 5.1 for text evaluation; Anaconda AI and ChatGPT5.1 for code evaluation. This work was supported by Australian Research Council Discovery Project DP220100040 (GPM), Australian Research Council Discovery Project DP250100345 (GPM), National Health and Medical Research Council Synergy Grant GNT2009851 (GPM)

## Methods

### Subjects

Male and female C57BL/6J mice (Australian Resources Centre; 8 - 10 weeks old) were housed in ventilation racks on corn cob bedding in a climate-controlled colony room under a 12:12 hr light/dark cycle. Mice had free access to food and water until 2 days prior to commencement of behavioural training. For the remainder of the experiment, they received 1 - 2 g of food and free access to water each day. Experiments were approved by the UNSW Animal Care and Ethics Committee and performed in accordance with the Animal Research Act 1985 (NSW), under both ARRIVE guidelines and the National Health and Medical Research Council Code for the Care and Use of Animals for Scientific Purposes in Australia (2013).

### Surgery

Mice were anaesthetized using 5% isoflurane in oxygen-enriched air, then fixed into a stereotax (Model 1900, Kopf Instruments). Mice were maintained on 2.5% isoflurane during surgery. 0.1 ml Marcaine (0.5%) was injected subcutaneously at the incision site. Ophthalmic gel (Viscotears, Alcon) was applied to avoid eye-drying. Mice received subcutaneous injections of antibiotic (Duplocillin, 0.15 ml/kg of body weight subcutaneously) and 5 mg/kg carprofen (Rimadyl, Zoetis) immediately post-surgery.

### Punishment task

All behavioral training and testing occurred in ENV-307A Standard Modular Chambers for Mouse (Med Associates). Each chamber was fitted with two retractable levers located either side of magazine for delivery of 20 mg grain pellets (BioServ, Dustless Precision Pellets). The chambers had a stainless-steel grid floor through which the punisher was delivered and were located in sound attenuating cabinets.

Mice first received 2 sessions of fixed-ratio (FR) 2 lever-press training. Both levers were extended, and every second press on either lever was reinforced with delivery of a pellet to an external magazine. Each lever retracted after being pressed 25 times. The session ended after 50 min or when both levers had retracted. Mice then received daily 40-min lever-press training sessions over the next 10 days. Left and right levers were alternately presented for 5 min. There were four trials per lever across 40 min. Lever-pressing was reinforced with pellet delivery on a variable-interval 30-s (VI30) schedule. The first lever to be extended (left or right) was fully randomised. Next, mice received punishment-training. Punishment was identical to the VI30 lever-press training sessions, except responses on one of the levers (designated punished lever) delivered a 0.5 s shock on a FR8 schedule independent of pellet outcome. The intensity of the punisher was 0.3mA except during calcium imaging (see below). The punished lever (left or right) was counterbalanced across animals. The other lever (designated unpunished lever) was rewarded by pellets but not punished by shocks.

The primary behavioral variable of interest was preference (suppression) ratio calculated as a/(a+b) where a = number of punished lever presses and b = number of unpunished lever presses. Values range from 0 to 1, where 0.5 indicates no lever preference and values < 0.5 indicate lower preference for punished lever relative to unpunished lever.

### Within-subjects yoking

After lever press training (see above), mice received a 40 min session of yoked training. Left and right levers were alternately presented for 5 min. There were four trials per lever across 40 min. The first lever to be extended (left or right) was fully randomised. This lever, designated ‘instrumental’, was rewarded on a VI30 schedule and punished (0.5s, 0.3mA) on an FR8 schedule. After 5-min, the instrumental lever was retracted and the second lever, designated ‘Pavlovian’, was inserted. This lever was also rewarded on a VI30 schedule and 0.5s, 0.3mA footshocks were delivered in a response non-contingent manner. Each mouse received the same number and timing of footshocks as it had received during its immediately preceding instrumental trial. Total lever presses in the final 5 min trial for each lever, latency to first lever press in the final 5 min trial for each lever, and suppression ratio for each lever, a/(a+b) where a = number of Trial 4 lever presses and b = number of Trial 4 punished lever presses + average number of presses on last day VI training. Values range from 0 to 1, where 0.5 indicates no change in lever preference and values < 0.5 indicate lower preference for punished lever relative to the end of VI training.

### Whole-brain Fos

After a single session of punishment (0.3mA, 0.5s footshock), mice were euthanized 45 min via an intraperitoneal injection of 0.1 mL sodium pentobarbitone (100 mg/kg). Mice were then transcardially perfused with 10–20 mL ice-cold saline (prewash), followed by 20 mL 4% paraformaldehyde (PFA) in phosphate-buffered saline (PBS). Brains were extracted and post-fixed in 4% PFA at 4 °C for 24 h, rinsed twice in PBS, and stored long-term in PBS containing 0.02% sodium azide at 4 °C until further processing.

Paraformaldehyde-fixed samples were preserved with using SHIELD reagents (LifeCanvas Technologies) using the manufacturer’s instructions. Samples were delipidated using LifeCanvas Technologies Clear+ delipidation reagents. Following delipidation samples were labelled using eFLASH (*69*) technology which integrates stochastic electrotransport (*70*) and SWITCH (*71*), using a SmartBatch+ (or SmartLabel) device (LifeCanvas Technologies). After immunolabeling (Primary: Rabbit anti-C Fos antibody, 3.5 ug, sample; Secondary: donkey anti-rabbit Setau-647, 5.25 ug per sample), samples were incubated in 50% EasyIndex (RI = 1.52, LifeCanvas Technologies) overnight at 37°C followed by 1 d incubation in 100% EasyIndex for refractive index matching. After index matching the samples were imaged using a SmartSPIM axially-swept light sheet microscope using a 3.6x objective (0.2 NA) (LifeCanvas Technologies).

Samples were registered to the Allen Brain Atlas (Allen Institute: https://portal.brain-map.org/) using an automated process (alignment performed by LifeCanvas Technologies). A propidium iodide channel for each brain was registered to an average atlas (generated by LCT using previously registered samples). Registration was performed using successive rigid, affine, and b-spline warping algorithms (SimpleElastix: https://simpleelastix.github.io/).

Automated cell detection was performed by LifeCanvas Technologies using a custom convolutional neural network created with the Tensorflow python package (Google). The cell detection was performed by two networks in sequence. First, a fully convolutional detection network (https://arxiv.org/abs/1605.06211v1) based on a U-Net architecture (https://arxiv.org/abs/1505.04597v1) was used to find possible positive locations. Second, a convolutional network using a ResNet architecture (https://arxiv.org/abs/1512.03385v1) was used to classify each location as positive or negative. Using the previously calculated Atlas Registration, each cell location was projected onto the Allen Brain Atlas in order to count the number of cells for each atlas-defined region.

Analyses were conducted in Python 3.10 using NetworkX (v2.8) and associated libraries (NumPy, Pandas, Matplotlib/Seaborn, python-louvain). For each group, Pearson correlation matrices of cell counts (Fos+ cells/mm^2^) across regions were computed and unweighted functional networks constructed by thresholding absolute correlations at r = 0.80 (diagonals set to zero) and binarizing edges.

Communities were identified with the Louvain algorithm (resolution γ = 1.0). To obtain robust partitions, we performed 200 independent runs with distinct random seeds (0–199) and formed a co-assignment matrix representing the probability that two nodes clustered together across runs. Edges with weight ≥ t = 0.60 were retained, and Louvain was re-run once on this thresholded graph to define consensus communities. Node stability was quantified as the proportion of runs in which each node’s label matched its consensus assignment. Partition similarity between groups was quantified using the Adjusted Rand Index (ARI) and Normalized Mutual Information (NMI) and a Jaccard similarity matrix was used to visualize cross-group module correspondence.

Node-level centrality included degree, betweenness, and closeness (*72–74*). To compare regional importance across groups, we Z-scored degree centrality values within each network and then computed the difference in Z-scores between Punishment and Control groups. This identified regions whose relative centrality increased or decreased under punishment. Network-level metrics included degree distributions, degree assortativity, clustering, characteristic path length, global efficiency, and diameter. Small-worldness (σ) was computed as σ = (C/C_rand)/(L/L_rand), using 200 degree-preserving rewired surrogates (seed = 42). Characteristic path length was evaluated on the largest connected component.

Robustness was assessed by targeted deletion: nodes were ranked by betweenness centrality and removed sequentially while tracking the largest connected component (LCC) fraction to yield percolation curves. To test the contribution of three hubs that increased in centrality under punishment (zona incerta; rostral linear nucleus raphe; posterior basolateral amygdala), we recalculated topology and global communicability after removing each hub individually and all three together. Communicability was defined as the sum of all pairwise values from the matrix exponential of the absolute adjacency (off-diagonal). Specificity was assessed by generating a null distribution of 5,000 random triple deletions (Punished network), from which a permutation p-value was computed.

### Chemogenetics

AAV5-hSyn-hM4D(Gi)-mCherry and AAV5-hSyn-mCherry were used. pAAV-hSyn-hM4D(Gi)-mCherry was a gift from Bryan Roth (Addgene viral prep # 50475-AAV5; http://n2t.net/addgene:50475; RRID:Addgene_50475). pAAV-hSyn-mCherry was a gift from Karl Deisseroth (Addgene viral prep # 114472-AAV5; http://n2t.net/addgene:114472; RRID:Addgene_114472). For triple silencing, 0.2 µl was microinjected into basolateral amygdala (coordinates in mm relative to bregma) (*75*) (−2.05 anteroposterior; ±3.30 mediolateral, and –5.0 dorsoventral), 0.1 µl into rostral zona incerta (rostral 1.0 anteroposterior: ±0.75 mediolateral, - 4.60 dorsoventral) and 0.15 µl into caudal zona incerta (−2.3 anteroposterior, ±1.4 mediolateral, and –4.50 dorsoventral). 0.15µl was injected into rostral linear nucleus raphe (coordinates relative to Lambda) (*75*) (±0.0 mediolateral, −0.65 anteroposterior and –4.05 dorsoventral). For single region silencing, the same coordinates and volumes were used. All infusions were made via 32-gauge conical-tipped microinfusion syringe (SGE Analytical Science) connected to a UMP3 with SYS4 Micro-controller microinjection system (World Precision Instruments) over a five-minute period. The needle remained in place for five minutes. Mice had at least 7 days recovery from surgery.

Behavioral training commenced no earlier than 14 days after surgery and followed the procedure described above. Mice received a dummy IP injection on the final day of VI lever press training to acclimatize them to injections. 30-min prior to the punishment session, all mice received an IP injection of CNO (3 mg/kg, 0.15ml, 0.5% v/v DMSO). 48 hr later they received a test of retention of punishment learning, identical to the VI30 training sessions. The firs 5-min block of each lever presentation was used as test data. Next, they received 2 - 3 days of VI30 lever-press training to equate responding on the levers. Then, mice received a 0.15ml IP vehicle injection 30 minutes prior to another punishment session which was otherwise identical to the first. Vehicle injections consisted of 0.5% v/v DMSO. Finally, mice received a second test session 48 hr later, identical to the first test session.

At the end of the experiment, mice were deeply anaesthetized via IP sodium pentobarbital (100 mg/kg) and perfused with 0.9% saline solution containing 1% sodium nitrite and heparin (5000 IU/ml), followed by phosphate buffer solution (0.1 M) with 4% paraformaldehyde. Brains were extracted, sliced coronally (40 mm) using a cryostat and processed for immunofluorescence.

### Spatial transcriptomics

After 10 days of VI lever press (see above), mice received a single 40-min session of punishment training (punishment (n = 4) or control (n = 4)). Immediately afterwards, mice were anaesthetized with isoflurane and then euthanized. Heads were submerged in ice-cold water and brains were rapidly extracted. Extracted tissue was immediately flash-frozen in liquid nitrogen and stored at – 80 °C until further processing. 14mm coronal sections were prepared using a cryostat and mounted onto a MERSCOPE slide (Lot #s 24060315, 24081215) and handled according to manufacturer guidelines (MERSCOPE™ User Guide - Fresh and Fixed Frozen Tissue Sample Preparation – 91600002 Rev B). Each slide contained 3 sections from a punished animal and 3 sections at the same coronal level from a control animal. Sections were chosen to span the anterior – posterior boundary of the target regions. Slides remained at −20°C for at least 5 min prior to 15 min incubation in fixation buffer at room temperature for 15 min. Slides were then washed 3 times with 5 mL 1X PBS (5 min each) before application of 5 mL 70% ethanol, and overnight storage at 4°C to permeabilize the tissue. Slides were stored at 4 °C prior to hybridization.

MERFISH using the MERSCOPE Gene Panel type - PanNeuro 500 gene mouse (Cat # 20300123) was performed according to manufacturer instructions, autofluorescence quenching was not used. Post sectioning the sections were kept at −20°C for 20min followed by 15min incubation in 4% PFA, washed 3 x 5min with RNAse free PBS and were kept in 70% ethanol to permeabilize the tissue. Following 5 days at 4°C in 70% v/v ethanol tissues were washed in sample prep wash buffer (Vizgen) and incubated in formamide buffer (Vizgen) at 37°C for 30min. After 50ml of the encoding probe mixture was applied directly to the tissue covered with parafilm and incubated for 48 hours at 37 °C in a humidified chamber. Following hybridization, samples were incubated in formamide buffer at 47°C for 30min, washed with sample prep wash buffer and incubated for 1 min in gel embedding solution (5 mL gel embedding premix (Vizgen), 25 µl 10% APS, 2.5 µl TEMED). Tissue was embedded in a thin layer of polyacrylamide gel using inverted embedding technique. Embedded tissues were then cleared in 5ml of clearing solution (5 mL clearing premix (Vizgen), 50 µl Proteinase K (NEB)) for up to 24 hours at 37 °C, until the tissue was optically transparent. Cleared samples were washed in sample prep wash buffer, stained with DAPI and PolyT (Vizgen) for 15min at RT and incubated in formamide for 10min prior to imaging. Imaging followed the Merscope Instrument Preparation guide (number 91500001) with 10mm section thickness selection and MERSCOPE 500gene imaging kit (10400006). For each slide, a 1 cm^2^ ROI was manually drawn based on DAPI and PolyT stains (10x magnification). Cell and transcript segmentation was completed via Vizgen Post-processing tool (VPT) using DAPI and PolyT - Cellpose model with Gaussian filter set to 30 and cell size 30. One midbrain slide was damaged during reactions, so n = 3 mice per group were used for midbrain analyses.

The MERFISH expression matrix was imported to Vizgen Merscope Visualizer and selected regions exported for further data analysis. First, cell-type annotation was performed using the Allen Institute’s MapMyCells tool. For each sample, the AnnData file from Vizgen viewer was uploaded to the MapMyCells web interface, to return a table containing, subclass_name, subclass_bootstrapping_probability, \\ supertype_name and supertype_bootstrapping probability. These annotations were imported into the MERSCOPE Visualizer to guide manual delineation of the amygdala, subthalamic–hypothalamic zone. Polygon coordinates and corresponding AnnData (.hdf5) files were exported, and all cells within the selected regions were extracted using custom Python scripts. The VTA region was automatically selected using a custom script that created a convex hull containing 97% of all cells (chosen by mahalanobis distance) with “VTA” in the subclass name. Similarly, the “RL” cells were selected by a similar method. These regions were manually verified.

Cells with total gene expression counts below the 5th percentile or above the 98th percentile for each slide were excluded. Expression values were normalized to a total count of 5,000 per cell, a threshold chosen to exceed the median total counts for the BLA while minimizing information loss. A UMAP plot of cells showed no visual clustering by slide and thus batch correction was not performed (**Fig S6**). The genes with the 100 lowest variances were calculated for each region and Anterior-Posterior combination and removed to prevent noise from very low variance genes affecting the results. The data were then passed into the cNMF package where gene expression was variance scaled so that genes on different expression scales would have equal weighting in the program calculation.

cNMF (*76*) was used to identify transcriptional programs. For each brain region and condition (control or punished), a principal component analysis scree plot showing log10 variance explained show an elbow around 10-19 programs for all regions. Accordingly, cNMF models were fit using 10–20 programs. Plots of reconstruction error and stability, together with principal component analysis scree plots showing log₁₀ variance explained, were used to select an optimal number of programs (n_programs) that maximized stability while minimizing error. The number of programs was kept constant between conditions when possible, except when this would cause a relatively unstable solution for a condition, in which case the number was allowed to differ by a maximum of 2 (although in practice control and punished never differed by more than one). When ambiguous, a slightly higher n_programs was chosen to ensure adequate resolution of transcriptional diversity.

For each region–condition combination, cNMF output included the top-weighted genes and cell-level program usages. Jaccard similarity matrices were computed to identify candidate punishment-specific programs. To assess robustness, leave-one-out resampling was performed by sequentially excluding each sample and recomputing programs for both conditions, yielding n program solutions per condition. Candidate punishment-specific programs were retained only if (i) no control leave-one-out run recapitulated them (Jaccard Index ≥ 0.5), and (ii) all punished leave-one-out runs did (Jaccard Index < 0.5 indicating a failure to reproduce). For the remaining punishment-specific programs, the dominant MapMyCells subclasses (those with usages in the top 30 of all subclasses) contributing to each program were identified and programs were selected only if at least 100 cells (regardless of usage) from one subclass were identified. We then identified all cells that used these programs as their top program and visualized program usage within each region of interest via spatial heatmaps.

### Calcium imaging

0.5ml AAV9-hSyn-GCaMP7f (pGP-AAV-syn-jGCaMP7f-WPRE was a gift from Douglas Kim & GENIE Project (Addgene viral prep # 104488-AAV; http://n2t.net/addgene:104488; RRID:Addgene 104488) was unilaterally injected using a 30-gauge conical-tipped microinfusion syringe (SGE Analytical Science) connected to a UMP3 with SYS4 Micro-controller microinjection system (World Precision Instruments). BLA coordinates were 1.5 mm posterior, ± 3.4 mm lateral and 4.8 mm ventral from bregma (*7*). The GRIN lens (0.5 mm diameter) was implanted 0.5 mm above. The lens was integrated into a baseplate that was secured to the skull using superglue and dental cement. Mice were allowed 6 weeks post-surgery recovery during which their condition was monitored daily (first week) then three times weekly (for the remainder of the experiment). Mice were single-housed post-surgery to prevent damage to GRIN lens implants. From 3 - 6 weeks post-surgery, mice were connected to the Miniscope weekly for habituation.

The multiday punishment task (*77*) commenced 6 – 7 weeks after surgery and first involved 10 days of lever press training on VI30 schedule as described above. These were followed by 6 days of 40-min punishment training where each session was as described above. To preserve opportunities to measure punished lever-press-related neural activity, shock intensity was gradually increased across punishment training (Days 1 - 2: 0.2 mA; Days 3 - 4: 0.3 mA; Days 5 - 7: 0.4 mA). There was a single 10-min choice session that involved concurrent presentations of both levers. During this choice session, lever presses were reinforced with pellet delivery on a unified VI60 schedule and there was no punishment. Finally, mice received 3 x 40-min punishment extinction sessions that were identical to initial lever press training as described above and no shock was delivered.

Mice were connected to the Miniscope daily throughout the experiment except on Days 3 - 5 of VI30 lever-press training, when mice were untethered to accelerate lever press acquisition. Recordings were made for key segments of each experiment session. On FR2 lever-press training days, recordings were made for the first and last 20 min of the session. On VI30 lever-press training, all punishment training days, and all punishment extinction days, recordings were made for the first and last 10 min of the session. Recordings were made for the entire 10 min of the choice test. Fluorescence Ca^2+^ imaging videos were acquired using nVista HD software (Inscopix Inc, Mountain View, CA, USA) with a field of view of approximately 600 x 600 mm, LED power of 0.3 - 0.5 mW/mm^2^, analogue gain of 1 - 3, and 2 x spatial down sampling. 12-bit images (1,000 x 1,000 pixels) were acquired at a frame rate of 30 Hz, with each pixel covering 0.6 x 0.6 mm in tissue. For each mouse, the same imaging parameters were used for all imaging sessions. The video data were streamed directly to a hard disk (90 −100 MB/s).

At the end of the experiment, mice were deeply anaesthetized via IP sodium pentobarbital (100 mg/kg) and perfused with 0.9% saline solution containing 1% sodium nitrite and heparin (5000 IU/ml), followed by phosphate buffer solution (0.1 M) with 4% paraformaldehyde. Brains were extracted, sliced coronally (40 mm) using a cryostat and processed for immunofluorescence to identify location of GRIN lens and GCaMP expression.

For each mouse, recordings from the same day were spliced into a single file and motion-corrected. Each image frame of the motion-corrected video was re-expressed in units of relative change in fluorescence, Δ*F*(*t*)/*F*_0_ = (*F*(*t*) – *F*_0_) /*F*_0_, where *F*_0_ is the mean image fluorescence obtained by averaging across the entire recording. This new video (Δ*F*/*F* video) allowed visualization of changes in cell fluorescence and thereby changes in their activity. Individual candidate BLA neurons were then identified via convex non-negative matrix factorization with spatiotemporal constraints (CNMFE). CNMFE parameters of cell diameter, minimum pixel correlation, and minimum peak-to-noise ratio were manually determined for each mouse to prevent over- or under-counting of cells. All other CNMFE parameters were set to default values. The full set of cellular activity transient data for each mouse on each day was then exported as a CSV file. Finally, for each mouse, a longitudinal registration algorithm pre-processed CNMFE cell sets from each session by normalizing images and creating a cell map. The algorithm then aligned each cell set to a reference and matched the cells of each aligned cell set to cells in previously matched sets. This created a new longitudinal cell set series that was saved as a CSV file for each mouse. There were 1,147 cells in total in this cell set from 6 mice. Mouse 1 – 6 = 139, 225, 196, 203, 111, 263 cells. Average number of sessions per cell was 3.12 sessions and subject range for this metric was 2.11 – 4.05.

MATLAB scripts were used to extract normalized and longitudinally registered Ca^2+^-dependent transients from CSV files for events categories of interest across the behavioral experiment. First, transients were low-pass (3 Hz) filtered to remove high-frequency noise identified via Fast Fourier Transform. Next, low-pass filtered transients were aligned with timestamps of events of interest recorded during behavioral sessions. These events were punished-lever-press, unpunished-lever-press, shock, and pellet consumption (hereafter: reward). Then, CNMFE-derived Δ*F t* /*F* values from −5 to +7 s around the following event categories were collated to generate peri-event transients: punished-lever-press alone (i.e., those not yielding shocks or rewards), unpunished-lever-press alone (i.e., those not yielding rewards), shock alone (i.e., those that did not co-occur with reward, although necessarily cooccurring with punished-lever-press) and reward alone (i.e., those that did not co-occur with shock). The −5 to +7 s peri-event interval was chosen to capture a sufficiently broad window of peri-event activity. Next, each peri-event transient was baselined (i.e., zeroed to average activity from −5 to −3 s before each event) then normalized by its sum squared deviation from zero (*78*).

K-means clustering was performed to categorize all cells co-registered across late lever-press training (days 9 - 10), early punishment (Day 1) and late punishment (Days 4 - 6) (n = 154 cells) according to their normalized peri-event transients. The validity of the 3-cluster solution was confirmed via 100% positive silhouette values, indicating each cell’s activity profile was well-represented by their cluster. Additional cells (those only partially co-registered across sessions) were incorporated into clusters if they had positive silhouette values for peri-event transients across shared sessions. Cluster 1 comprised 165 cells, Cluster 2 comprised 270 cells, and Cluster 3 comprised 176 cells. Within each cluster, normalized peri-event transients of all neurons were averaged into cluster-based kernels for each event category of interest at each key experiment phase (e.g., the Cluster 1 peri-reward early-punishment kernel represents the averaged normalized activity of Cluster 1 neurons detected around all reward events during early punishment). Significant cluster-based kernel transients for the −5 to +5 s peri-event period were identified via 95% bootstrapped confidence intervals (10).

Within each cluster, similarity between punished-lever-press, unpunished-lever-press, shock, and reward activity kernels was quantified by deriving a normalized fit score for each pairwise kernel comparison. The normalized fit score (−1 to 1) was calculated as the dot product of two peri-event kernel transient vectors (each normalized according to its sum square deviation from zero). A positive fit score indicates that two kernel transients deviate from baseline in a similar way across the peri-event interval. A negative fit score indicates that two kernel transients deviate from baseline in inverse ways. Zero indicates that two kernel transient vectors are orthogonal to one another.

To determine whether peri-event kernel transients across sessions were significantly similar (fit > 0) or inverse (fit < 0) to each other, 95% CI limits for fits were obtained from 2.5 and 97.5 percentiles of bootstrapped fit distributions (fits of randomly resampled mean transients, 1000 iterations). Two transients were deemed significantly similar if the fit CI was entirely above zero, or significantly inverse if the fit CI was entirely below zero. To visualize similarity and dissimilarity between each pair of cluster-based kernels, kernel activity similarity matrices (heat maps) showing all fit scores and results of significance tests were generated. To visualize the overall similarity/dissimilarity of peri-event activity kernels across sessions, fit scores were converted into fit distances (1 - fit score), such that identical kernels (fit score = 1) would have fit distance = 0, and mirror-image inverse kernels (fit score = −1) would have fit distance = −2. Kernel coordinates in 2D space were obtained via multidimensional scaling (MATLAB mdscale function, criterion = metrics-stress) using fit distances as input.

To examine the relationship between punished-lever-press–related BLA neuronal activity and punishment avoidance, we computed cluster-based kernel relative fit scores and correlated them with punished-response suppression ratios for each mouse across five experimental phases: lever-press training (days 9–10), early punishment (day 1), late punishment (days 4–6), early extinction (day 1), and late extinction (day 3). Relative fit scores quantified the similarity of each mouse’s average punished-lever-press-related BLA activity at each phase relative to pre- and late-punishment. For each phase, the relative fit score was defined as the normalized fit of the phase-specific punished-lever-press kernel to the mouse’s late-punishment (days 4–6) kernel, minus its normalized fit to the pre-punishment (training days 9–10) kernel. Positive values indicate that punished-lever-press-related activity more closely resembled late-punishment than pre-punishment patterns, whereas negative values indicate the reverse. Punished-response suppression ratios provided an index of punishment avoidance by quantifying the self-normalized change in punished-lever-press rate during each phase relative to pre-punishment. Suppression ratios were calculated as: Pun LP rate/ (Pun LP rate + Unpun LP rate). Values range from 0 to 1, where 0.5 indicates no change relative to pre-punishment, values > 0.5 indicate higher punished-lever-press rates, and values < 0.5 indicate lower rates. Relationships between relative fit scores and suppression ratios were assessed using linear regression.

---

**Fig S1.**
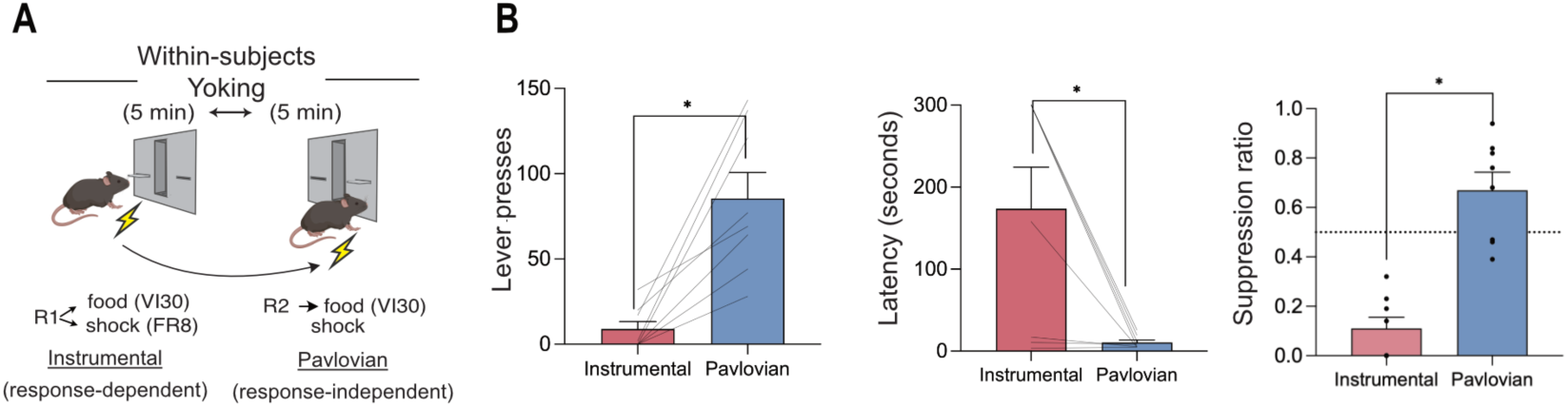
Within-subjects yoking: (A) Schematic of the within-subjects yoking procedure. Mice (N = 8) received alternating 5-min access to two levers. On the instrumental lever (R1), food was delivered on a VI30 schedule and footshock on an FR8 schedule. For each mouse, shock delivery times from R1 were recorded and replayed in a response-independent manner during access to the Pavlovian lever (R2), which delivered food on a VI30 schedule. This design equates Pavlovian contingencies across R1 and R2 while preserving the instrumental contingency on R1. (B) Behavioral measures. Trial 4 lever pressing was selectively suppressed on the instrumental lever (left), accompanied by increased response latency (middle) and reduced suppression ratios (computed relative to last day of VI training) (right). Statistical analysis: p < 0.05, dependent means t-test (D).

**Fig S2.**
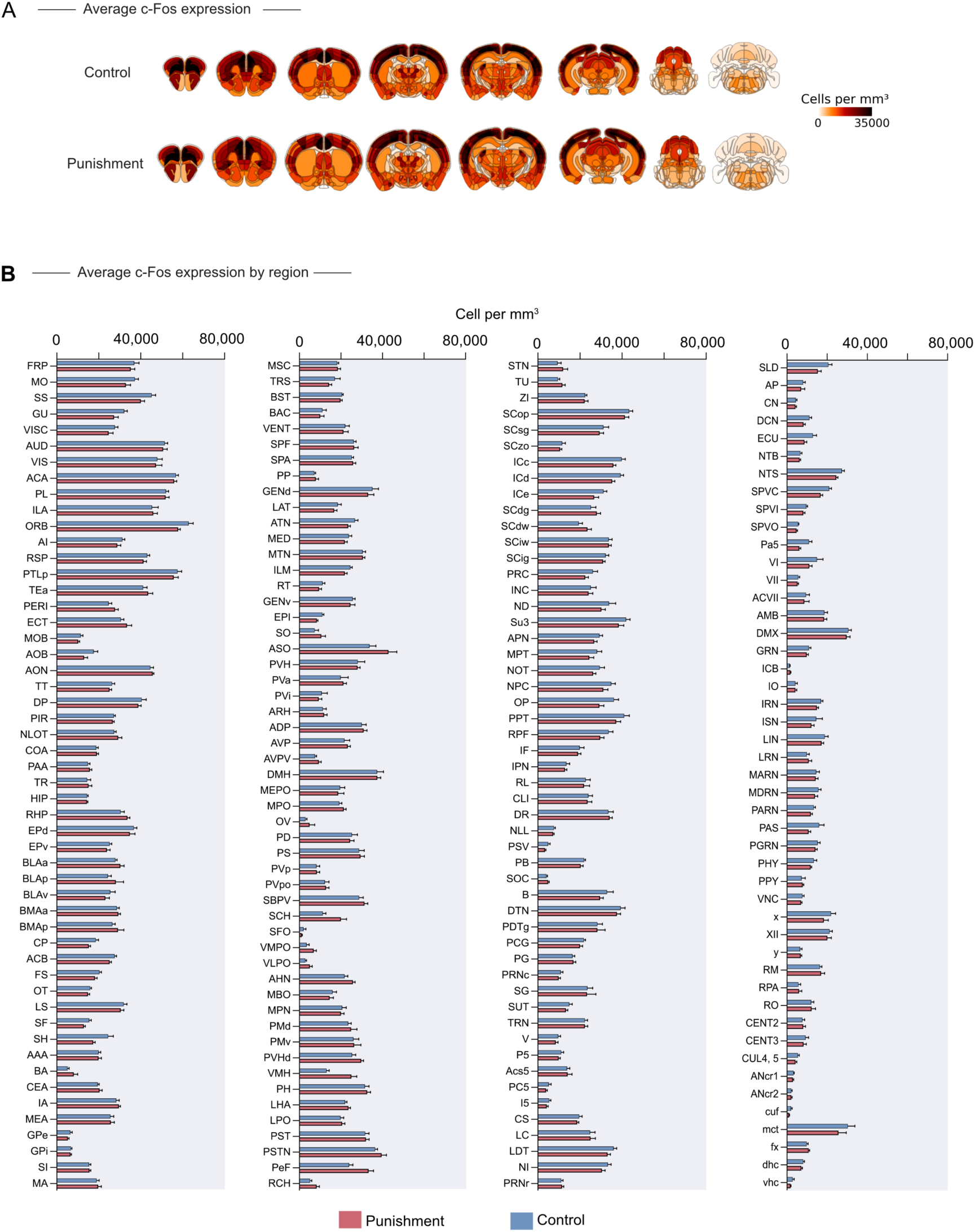
Whole brain c-Fos expression. (A) representative coronal brain sections showing averaged c-Fos+ cells per mm^2^. (B) Mean and SEM c-Fos+ cells per mm^2^ for Control and Punishment group across 206 brain regions.

**Fig S3.**
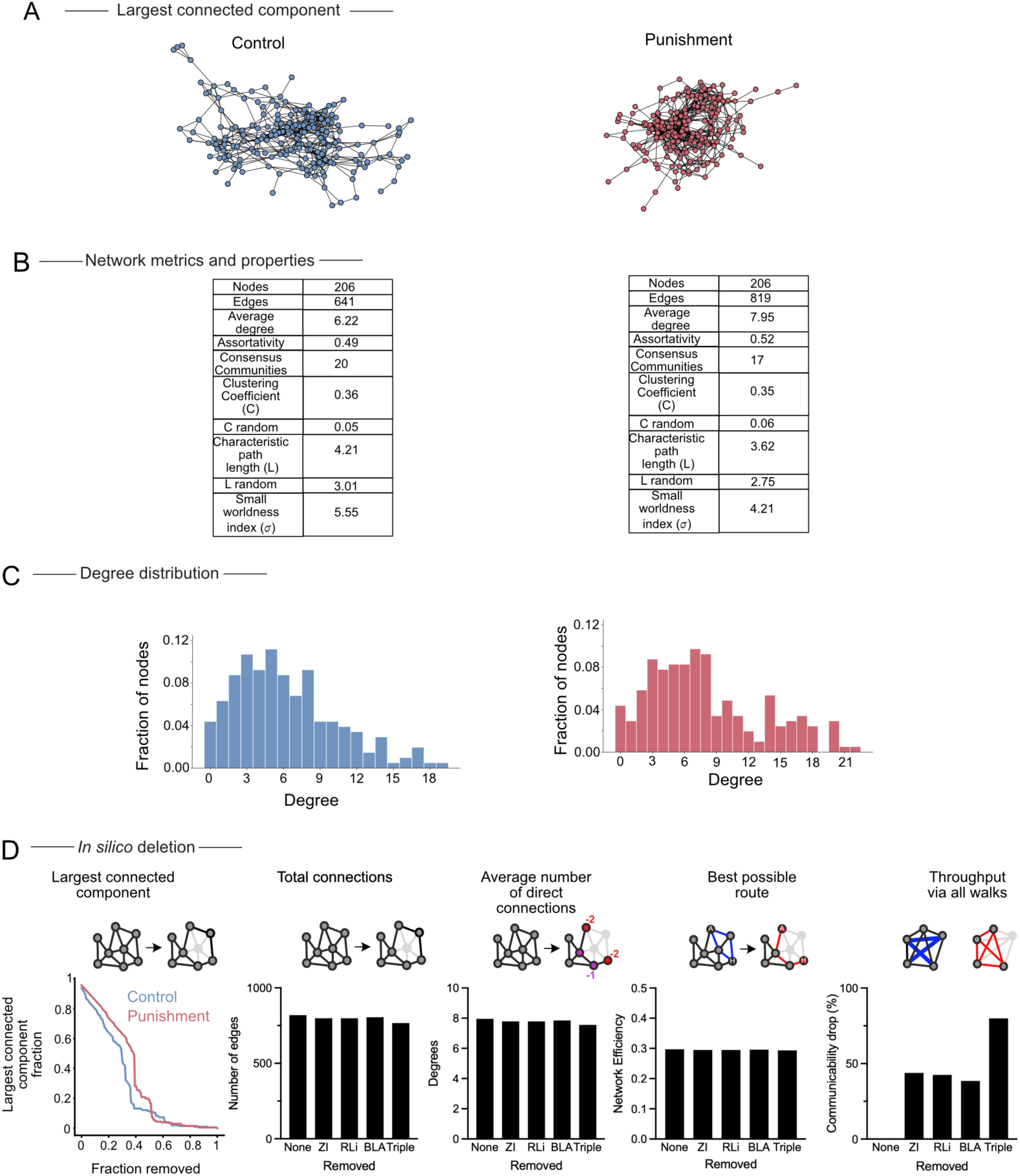
Network analyses of whole brain c-Fos expression. (A) Network structure for Control and Punishment groups. (B) Key network properties for Control and Punishment groups. (C) Frequency distribution of degree centrality counts for Control and Punishment groups. (D) Percolation curve and impact of individual or triple node deletions on network metrics.

**Fig S4.**
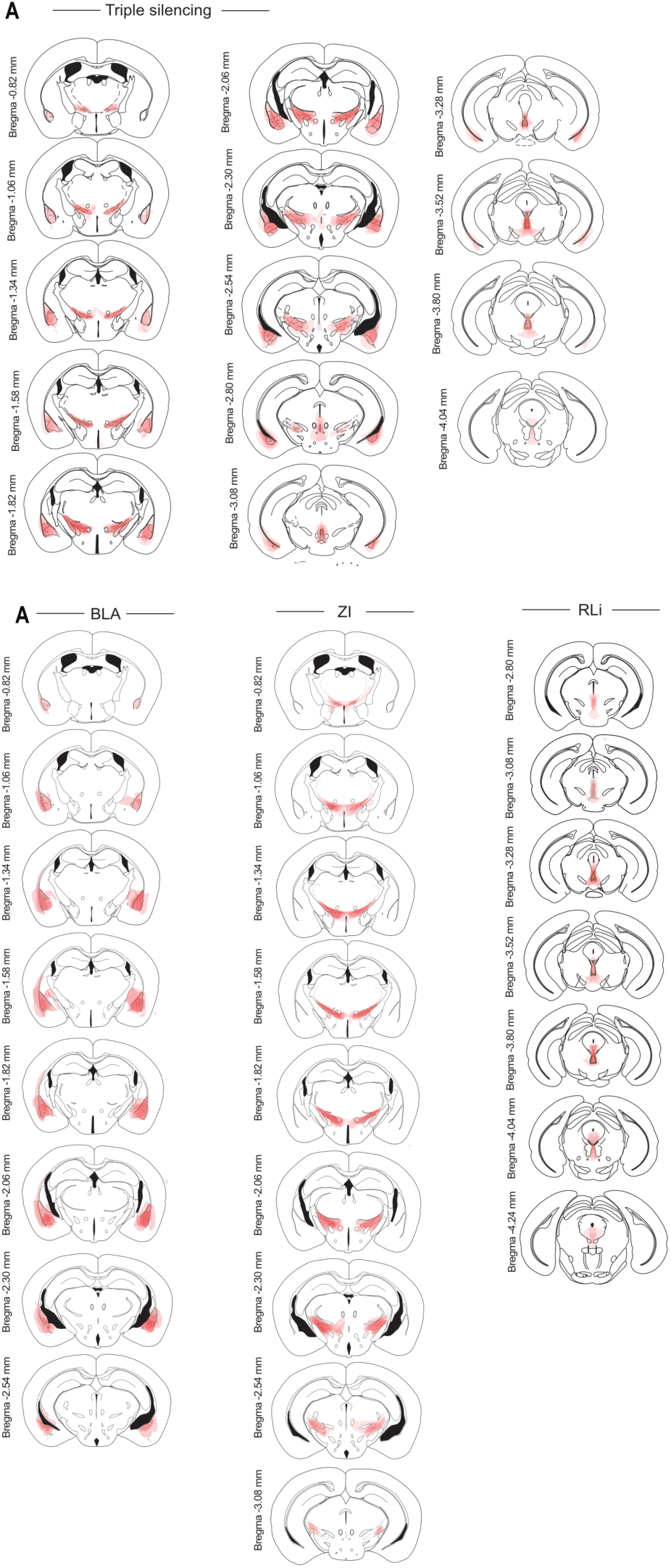
Histological verification of hM4Di expression. Location of hM4Di expression in amygdala, subthalamic–hypothalamic zone, and ventral midbrain tegmentum on Paxinos and Franklin atlas.

**Fig S5.**
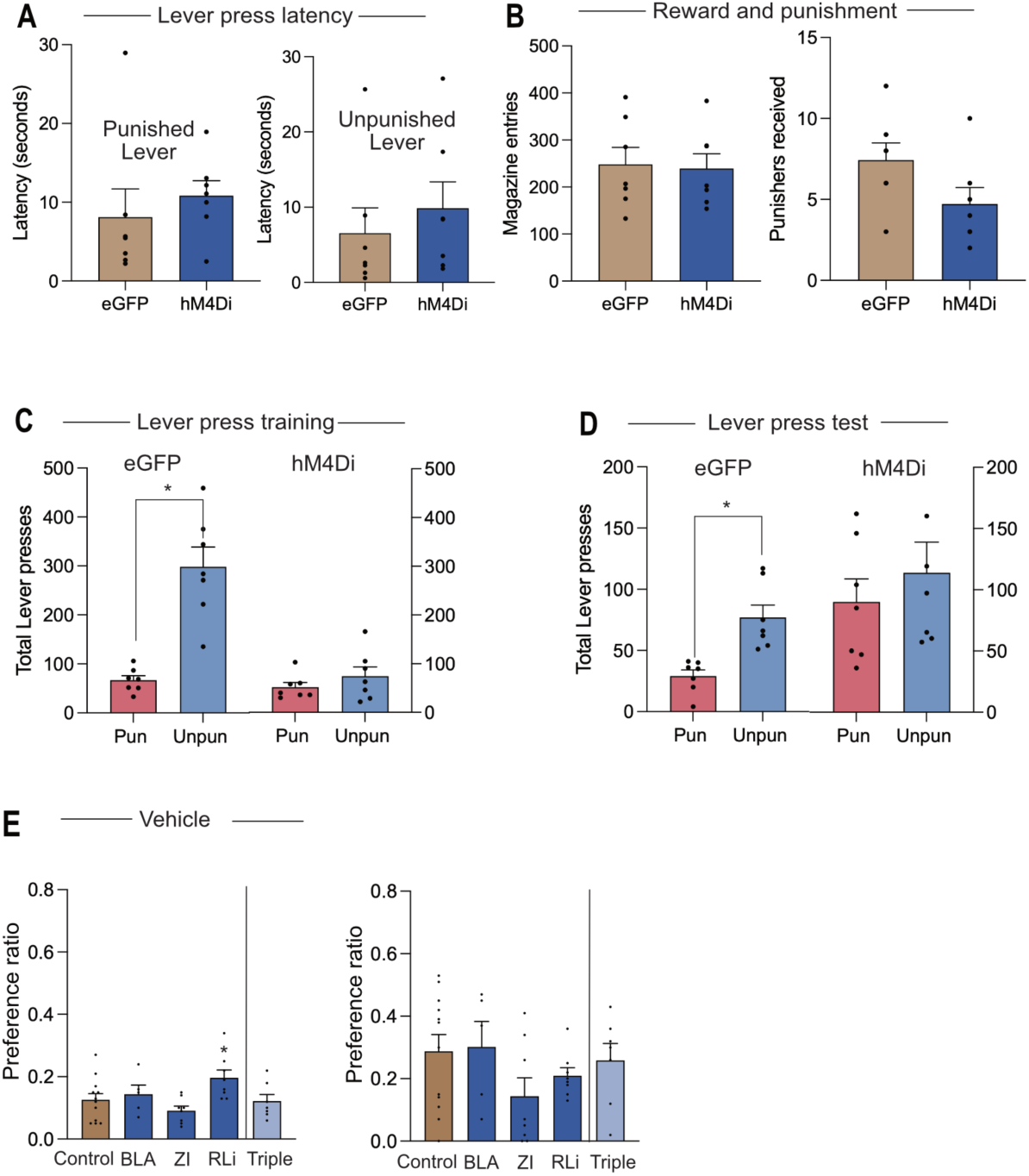
Chemogenetic silencing. (A) Latencies to first punished and unpunished lever press. (B) Total magazine entries and total punishers received. (C) Total punished and unpunished lever presses during 40 minute training. (D) Total punished and unpunished lever presses during 5 minute test. (E) Training and test preference ratios for vehicle condition. Statistical analysis: * = p < 0.05, was assessed by independent means t-test (A,B) dependent means t-test (C, D), one-way analysis of variance (ANOVA) and follow-up pairwise comparisons against Control (E). All data are presented as mean ± SEM.

**Fig S6.**
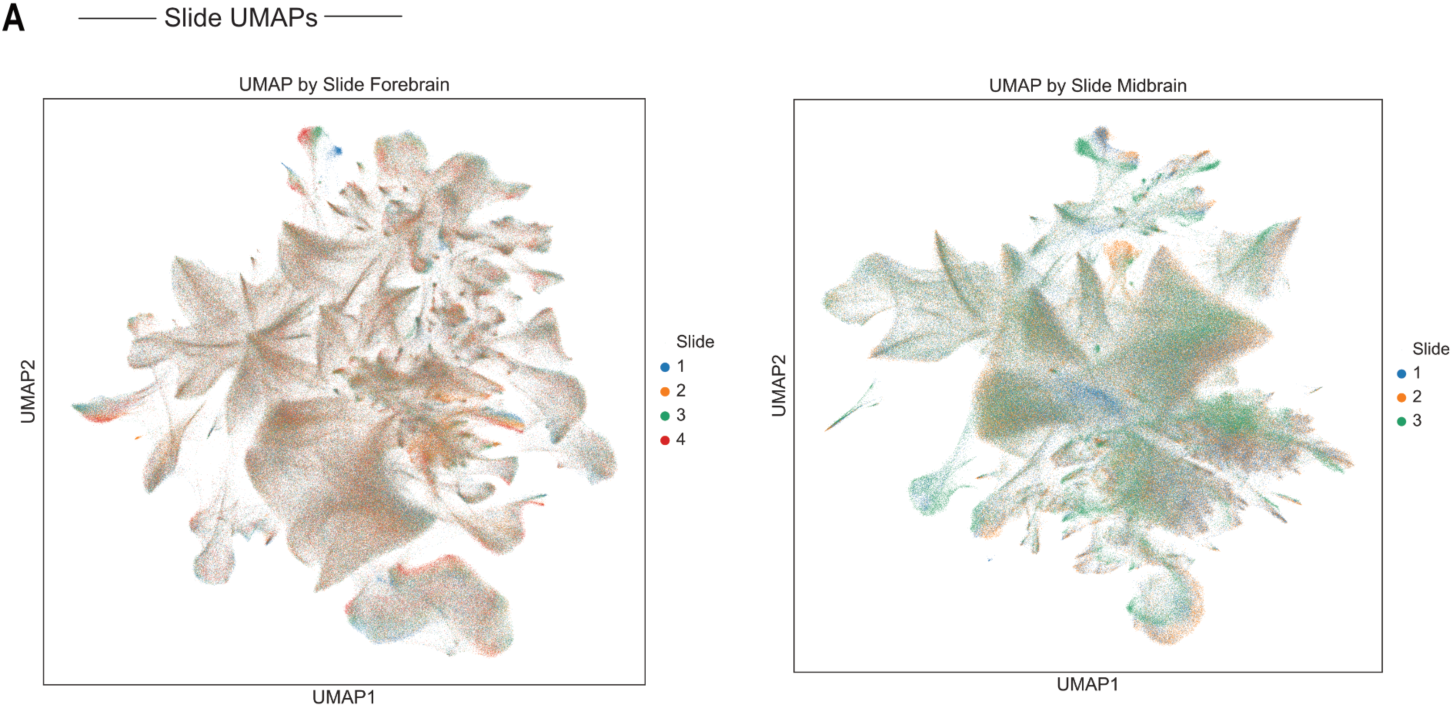
UMAP plots for all cells in forebrain and midbrain slides. Each dot represents a cell and the colors represent different slides. There was no visual clustering of the cells by slide. The UMAPs were generated to 40 components using default parameters and the first 2 components were visualised.

**Fig S7.**
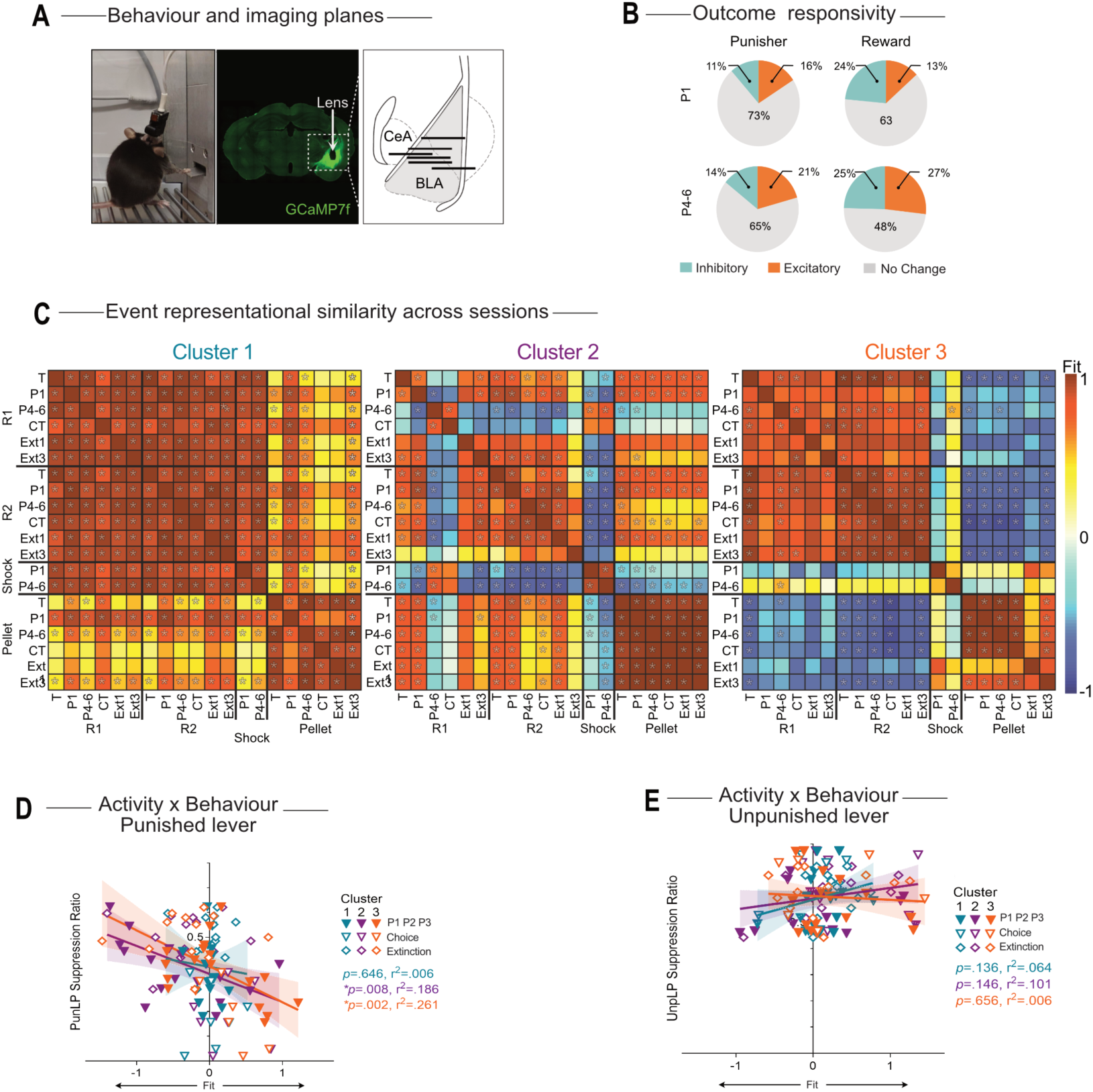
Basolateral amygdala ensembles encode outcome-specific representations and link to behavioral suppression. (A) Behavior and imaging planes. Example lever press during imaging (left), representative GCaMP7f expression and GRIN lens placement (middle), location of imaging planes (right). (B) Outcome responsivity. Proportion of neurons showing excitation, inhibition, or no change to punisher (N = 514) and reward (N = 573) outcomes during early (P1) and late (P4–6) punishment sessions. (C) Event representational similarity across sessions. Kernel activity similarity matrices (heat maps) showing fit scores and significance outcomes for all pairwise comparisons between cluster-based kernels. (D–E) Activity × behavior. Correlations between ensemble fit (mean normalized kernel fit for punished or unpunished lever events) and behavioral suppression ratios for the punished (D) and unpunished (E) levers. Lines show linear fits ± 95% CI. Statistical analysis: * = p < 0.05. Chi-square (B), 95% CI limits for fit values were obtained from the 2.5 and 97.5 percentiles of bootstrapped fit distributions (random resampling of mean transients, 1000 iterations) (C), linear regression (D, E).

